# Replicating RNA platform enables rapid response to the SARS-CoV-2 Omicron variant and elicits enhanced protection in naïve hamsters compared to ancestral vaccine

**DOI:** 10.1101/2022.01.31.478520

**Authors:** David W. Hawman, Kimberly Meade-White, Chad Clancy, Jacob Archer, Troy Hinkley, Shanna S. Leventhal, Deepashri Rao, Allie Stamper, Matthew Lewis, Rebecca Rosenke, Kyle Krieger, Samantha Randall, Amit P. Khandhar, Linhue Hao, Tien-Ying Hsiang, Alexander L. Greninger, Michael Gale, Peter Berglund, Deborah Heydenburg Fuller, Kyle Rosenke, Heinz Feldmann, Jesse H. Erasmus

## Abstract

In late 2021, the SARS-CoV-2 Omicron (B.1.1.529) variant of concern (VoC) was reported with many mutations in the viral spike protein that were predicted to enhance transmissibility and allow viral escape of neutralizing antibodies. Within weeks of the first report of B.1.1.529, this VoC has rapidly spread throughout the world, replacing previously circulating strains of SARS-CoV-2 and leading to a resurgence in COVID-19 cases even in populations with high levels of vaccine- and infection-induced immunity. Initial studies have shown that B.1.1.529 is less sensitive to protective antibody conferred by previous infections and vaccines developed against earlier lineages of SARS-CoV-2. The ability of B.1.1.529 to spread even among vaccinated populations has led to a global public health demand for updated vaccines that can confer protection against B.1.1.529. We report here the rapid development of a replicating RNA vaccine expressing the B.1.1.529 spike and show that this B.1.1.529-targeted vaccine is immunogenic in mice and hamsters. Interestingly, we found that mice previously immunized with A.1-specific vaccines failed to elevate neutralizing antibody titers against B.1.1.529 following B.1.1.529-targeted boosting, suggesting pre-existing immunity may impact the efficacy of B.1.1.529-targeted boosters. Furthermore, we found that our B.1.1.529-targeted vaccine provides superior protection compared to the ancestral A.1-targeted vaccine in hamsters challenged with the B.1.1.529 VoC after a single dose of each vaccine.

**One Sentence Summary:** Rapidly developed RNA vaccine protects against SARS-CoV-2 Omicron variant

## Introduction

Since emerging in late 2019 in China, severe acute respiratory disease - coronavirus 2 (SARS-CoV-2) has caused hundreds of millions of infections and the associated disease, COVID-19, resulting in millions of deaths. As of early 2022, the virus continues to circulate throughout the world. To address the pandemic, multiple vaccines were rapidly developed and early reports showed high levels of protection against symptomatic infection, severe disease and hospitalization (*1*). However, continued viral evolution has given rise to variants of concern (VoC) with abilities to evade vaccine- or infection-induced immunity and increase transmissibility (*2–5*). However, the recent emergence of the B.1.1.529 (Omicron) VoC in late 2021 (*6*) has resulted in an unprecedented resurgence of COVID-19 cases with many countries reporting record case numbers. Remarkably, in the United States, the B.1.1.529 VoC represented less than 1% of cases in early December 2021 but by mid-January 2022, was responsible for >99% of cases (*7*). The remarkable replacement of previously circulating SARS-CoV-2 strains by B.1.1.529 is likely due to 1) the ability of the B.1.1.529 to evade either vaccine- or infection-induced immunity (*8-11*) enabling B.1.1.529 to spread among previously resistant populations, and 2) increased transmissibility as many of the described mutations have been previously implicated in enhancing the receptor binding domain’s affinity for the ACE2 receptor (*12, 13*) with B.1.1.529 also appearing to replicate efficiently in the upper respiratory tract promoting efficient transmission in rodent models of infection (*14*). The emergence of B.1.1.529 and its resistance to previously acquired immunity has resulted in a public health demand for updated vaccines that can limit infection and transmission of the B.1.1.529 VoC to address the ongoing public health threat posed by SARS-CoV-2. Genetic immunization-based vaccine technologies, including those based on mRNA modalities, provide the ability to rapidly respond to such changes in the virus and the currently approved mRNA vaccines for COVID-19 are in the process of being updated (*15*).

Previously, we reported on the development of a cationic nanocarrier, Lipid InOrganic Nanoparticle (LION), for delivery of a replicating RNA (repRNA) encoding the ancestral Spike of SARS-CoV-2 (*16*) as well as those of the B.1.1.7 and B.1.351 lineage viruses (*17*). Investigatory COVID-19 vaccine products based on the LION/repRNA platform are currently under evaluation in an ongoing phase II/III trial in India, under the drug product designation HDT/Gennova COVID-19 (HGC019, clinical trial identifier CTRI/2021/09/036379), and in ongoing phase I trials in South Korea under the name QTP104, as well as in Brazil and the US under the name HDT-301. In contrast to lipid nanoparticle (LNP)-based approaches for RNA delivery, cationic nanocarriers provide a distinct stability and independent manufacturing advantage that enables their stockpiling for rapid response to emerging diseases, including variants of SARS-CoV-2. Upon manufacture of updated repRNAs, the two components are simply combined and mixed by inversion, prior to loading of syringes for intramuscular injection. We report here the rapid development of a B.1.1.529-targeted repRNA and demonstrate that this vaccine is immunogenic and confers improved protection against B.1.1.529 infection in a hamster model compared to the ancestral A.1-specific vaccine, indicating that B.1.1.529-targeted vaccines are likely needed for optimal protection against B.1.1.529 infection. Importantly however, we also evaluated the immunogenicity of a B.1.1.529-targeted vaccine as a booster in A.1-pre-immune mice and found that pre-existing immunity could negatively impact VoC-targeted booster responses.

## Results

### Design and production of a B.1.1.529-targeted repRNA-CoV2S

Following the November 25^th^, 2021, announcement of a new VoC, first discovered in South Africa and designated by the WHO as the Omicron variant, we initiated nonclinical activities to rapidly update our repRNA-CoV2S, including vaccine design, as well as *in vitro* and *in vivo* evaluations, to inform ongoing clinical studies of our COVID-19 vaccine (Fig. 1A). We screened the sequences deposited on GISAID available at that time for complete coverage of the spike (S) open reading frame (ORF) andselected EPI_ISL_6699769, deposited by de Oliveira *et al* (*6*), to design an updated S ORF for insertion into our previously described repRNA-CoV2S (*16, 17*) (Fig. 1B). Following the rapid synthesis of three overlapping tiles, we cloned and sequenced-verified B.1.1.529-repRNA-CoV2S prior to production of RNA for *in vitro* qualification using an *in vitro* potency assay. A half-maximal effective concentration (EC_50_) of 13.3 ng/well was measured (Fig. 1C), an expected potency within the range of previous repRNA-CoV2 drug substances (*17*).

**Figure 1.**
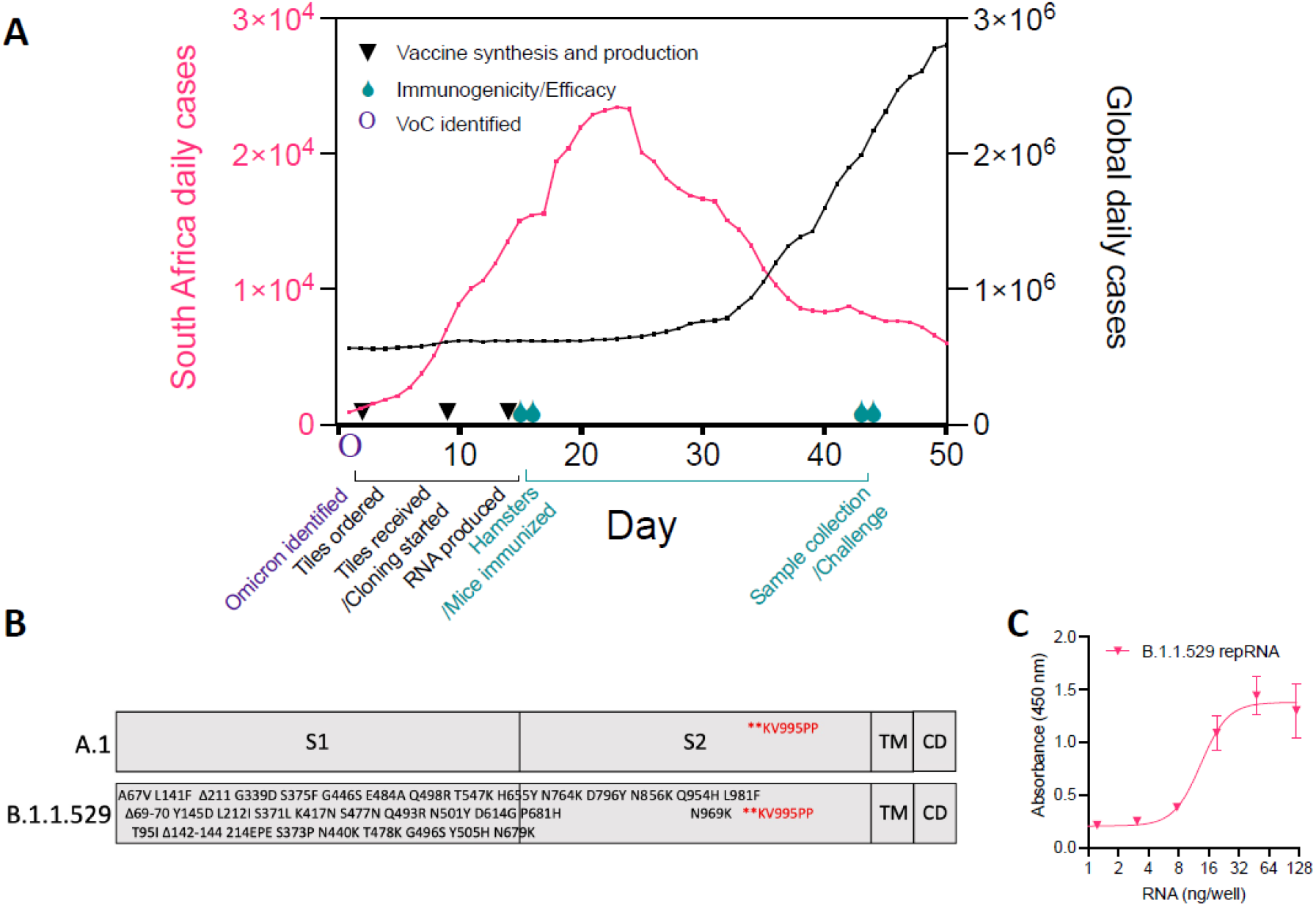
Design, production, and testing of a B.1.1.529-targeted repRNA-CoV2S vaccine in response to the Omicron wave. (**A**) Timeline of nonclinical studies to evaluate an updated B.1.1.529-targeted repRNA-CoV2S in the context of the ongoing Omicron wave (data sourced from ourworldindata.org). (**B**) Design of A.1 and B.1.1.529 spike (S) open reading frames expressed from repRNA, including the amino acid substitutions, insertions, and deletions present in GISAID accession no. EPI_ISL_6699769, relative to the A.1 S and their respective locations within the S1, S2, transmembrane (TM), and cytoplasmic (CD) domains. (**C**) *in vitro* potency assay of B.1.1.529-repRNA-CoV2S in baby hamster kidney (BHK) cells.

### Vaccination of pre-immune or naïve animals elicits differential antibody responses

Due to the prevalence of SARS-CoV2 immunity in the global population and the current need to boost pre-immune individuals we initiated a single or two-dose booster study in three groups of pre-immune mice. These animals had previously received 1μg doses, spaced 28 days apart, of either a prime/boost A.1-repRNA-CoV2S vaccine (A.1 Spike 2X), or a prime-only A.1-repRNA-CoV2S followed by a control boost of an influenza HA-repRNA (A.1 Spike 1X), or a control prime/boost of an influenza HA-repRNA vaccine (Influenza HA 2X) (Fig. 2A). The inclusion of the HA-repRNA vaccinations were important to control for immune response to the repRNA backbone that encodes the nonstructural proteins of Venezuelan equine encephalitis virus. At 24 days after their second vaccination, all animals received two 1μg booster doses of B.1.1.529-repRNA-CoV2S on days 0 and 28 (Fig. 2A) Sera was collected on on days 28 and 42 to evaluate antibody responses after each booster, by enzyme-linked immunosorbent assay (ELISA) measured against recombinant A.1 spike, or the A.1 or B.1.1.529 receptor binding domains (RBDs) (Fig. 2B), and by live-virus 80% plaque reduction neutralization test (PRNT_80_) measured against live B.1.1.529 virus (Fig. 2C). After the first and second B.1.1.529-repRNA-CoV2S boosters, we observed no significant changes in bAb specificity or magnitude against any of the three recombinant proteins the A.1 Spike 2X pre-immune animals, whereas bAb responses against A.1 spike and B.1.1.529 RBD appeared to increase after each B.1.1.529 booster in the A.1 Spike 1X pre-immune animals, although these changes were not significant (Fig. 2B). In contrast, the “naïve” influenza HA 2X pre-immune animals that received 1 or 2 boosters of B.1.1.529-repRNA-CoV2S exhibited significant increases in A.1 spike- and B.1.1.529 RBD-bAb responses. These increased from undetectable pre-boost to 28 and 23 μg/ml (geometric means), respectively, post-boost, but no significant changes in A.1 RBD-bAb responses (Fig. 2B). Interestingly, the second booster did not appear to offer any benefit in the influenza HA 2X pre-immune mice, and only provided a 4-fold boost (from 3 to 12.8 μg/ml in geometric means) in the A.1 spike 1X pre-immune animals against B.1.1.529 RBD. In terms of B.1.1.529-specific neutralizing antibody titers (nAbs), a single B.1.1.529-repRNA-CoV2S boost failed to induce detectable responses in A. 1 Spike 2X pre-immune animals, and a detectable 1:20 PRNT_80_ titer in 1 of the 5 A.1 Spike 1X pre-immune animals (Fig. 2C). However, in the “naïve” influenza HA 2X pre-immune animals, a single B. 1.1.529-repRNA-CoV2S booster elicited a detectable response in 3 out of 5 animals with a geometric mean titer (GMT) of 1:40 and 1:101 in the three responders (Fig. 2C).

**Figure 2.**
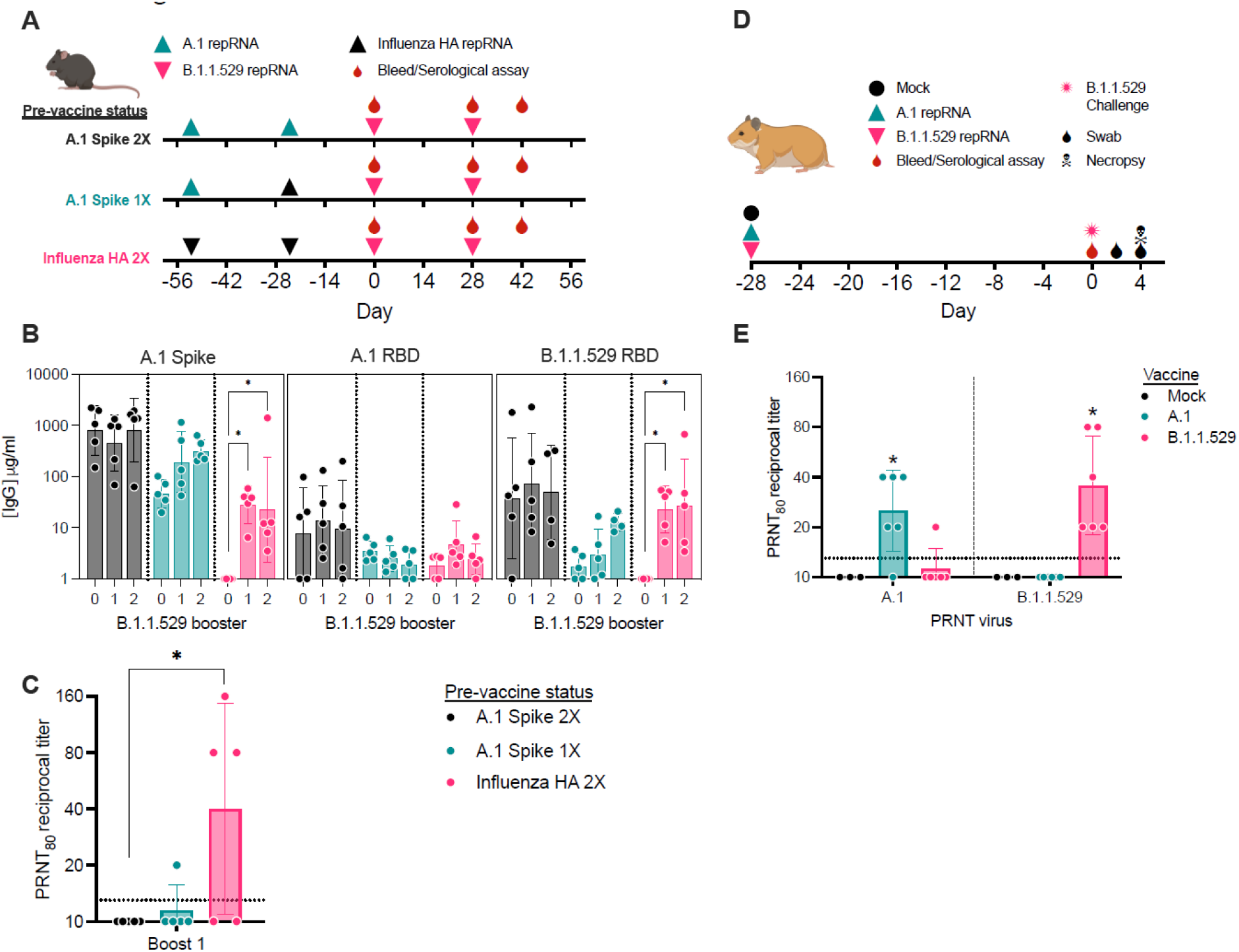
Antibody responses in pre-immune mice or naïve hamsters. (**A**) Study design for evaluation of B.1.1.529-repRNA-CoV2S immunogenicity in pre-immune mice. C57BL/6 mice (n=5/group) received 1μg doses in either a prime/boost of A.1-repRNA-CoV2S (A.1 Spike 2X), a prime with A.1-repRNA-CoV2S and boost with influenza HA-repRNA (A.1 Spike 1X), or a prime/boost with influenza HA-repRNA (Influenza HA 2X) prior to all groups receiving a 1μg boost on day 0, followed by a bleed and second boost on day 28 with a final bleed on day 42. (**B**) Day 0, 28, and 42 sera were evaluated for binding antibody responses to recombinant A.1 Spike, A.1 receptor binding domain (RBD), or B.1.1.529 RBD by enzyme linked immunosorbent assay (ELISA). (**C**) Day 28 and 42 sera were evaluated for neutralizing antibody responses against B.1.1.529 virus by 80% plaque reduction neutralization test (PRNT_80_). (**D**) Study design for evaluation of comparative immunogenicity and efficacy of a single-dose A.1- or B.1.1.529-repRNA-CoV2S vaccines against B.1.1.529 challenge in Syrian hamsters. Syrian hamsters (n=6/group) were mock vaccinated with saline or with 20μg of either A.1-repRNA-CoV2S or B.1.1.529-repRNA-CoV2S on day −28. Then on day 0, all animals were bled followed by an intranasal challenge with 1000 tissue culture 50% infectious doses (TCID_50_) and swabs collected on days 2 and 4 prior to necropsy on day 4 when blood and tissue samples were collected. (**E**) Day 0 sera was then assayed for A.1 or B.1.1.529-targeted neutralization activity by PRNT_80_. Indicated statistical comparisons performed using one-way ANOVA of log transformed values with Dunnett’s multiple comparison test. *p<0.05. Comparisons without indicated p-values were non-significant (p>0.05)

We previously reported on variant-specific immunogenicity of a 20μg prime/boost of A.1-, B.1.1.7-, or B.1.351-specific repRNA-CoV2S in naïve Syrian hamsters (*17*). However, given the need to generate rapid immunogenicity and efficacy data to inform manufacturing and clinical development activities of an updated vaccine, we opted to evaluate single-dose comparative immunogenicity/efficacy in the same naïve hamster model in parallel to the mouse study described above. Here, we vaccinated naïve hamsters with a single 20 μg dose of either A.1- or B.1.1.529-repRNA-CoV2S followed by B.1.1.529 challenge 28 days later (Fig. 2D). In contrast to the B.1.1.529-repRNA-CoV2S responses in pre-immune mice, a single 20μg dose of either an A.1- or a B.1.1.529-repRNA-CoV2S in naïve hamsters was able to elicit homologous A.1- or B.1.1.529-targeted nAb responses (Fig 2E), with GMTs of 1:25 or 1:36, respectively. However, heterologous nAb responses were mostly undetectable with only 1 animal, out of 6, that received a B.1.1.529-repRNA-CoV2S vaccination exhibiting a 1:20 PRNT_80_ titer against A.1 virus. All 6 A.1-repRNA-CoV2S vaccinated hamsters exhibited undetectable PRNT_80_ titers against B.1.1.529 virus, 28 days after a single vaccination.

### Single dose of B.1.1.529-targeted vaccine in naïve hamsters confers significant protection against B.1.1.529 challenge

Our hamster sera neutralization data showed that the A.1 vaccine did not elicit antibodies able to neutralize the B.1.1.529 variant suggesting that A.1-repRNA-CoV2S would provide incomplete protection against the B.1.1.529 variant. To test this hypothesis, we challenged the hamsters with 1000 TCID_50_ of B.1.1.529 via the intranasal route. On days 2 and 4 post infection (PI), oral swabs were collected and on day 4 animals were euthanized for evaluation of viral loads in the respiratory tract. Consistent with previous reports on mild clinical disease in hamsters infected with B.1.1.529 (*18–20*) we observed no overt clinical disease in any groups after challenge as evidenced by no weight loss in any group (Supplemental Figure 1). On day 2 PI, neither A.1-nor B.1.1.529-repRNA-CoV2S vaccinated animals had reduced amounts of viral RNA compared to mock-vaccinated animals (Figure 3A). However, by day 4 PI, both A.1- and B.1.1.529-repRNA-CoV2S vaccinated animals had significantly reduced amounts of viral RNA in the oral swabs (Figure 3A) suggesting that vaccination with either the A.1- or B.1.1.529-repRNA-CoV2S could reduce the duration of viral shedding. However, we were largely unable to detect infectious virus within the oral swabs at any timepoint (Figure 3B). Within the respiratory tract on day 4 PI, we found that A.1-repRNA-CoV2S vaccination significantly reduced viral RNA and infectious virus burden in the nasal turbinates but did not significantly reduce viral loads in the trachea (Figure 3C&D). In contrast, B.1.1.529-repRNA-CoV2S vaccination led to significantly reduced viral RNA burden in all three tissues (Figure 3C) and infectious virus was only detected in the lung tissue of one hamster while no infectious virus was detected in the trachea or nasal turbinates of any B.1.1.529-repRNA-CoV2S vaccinated animals (Figure 3D). Overall, infectious virus was rarely detected in the lungs of any animal. Instead, infectious virus was detected mainly in the upper respiratory tract (Figure 3D), a result that is consistent with the absence of clinical disease in the B.1.1.529 infected hamsters. These data show that the A.1-repRNA-CoV2S vaccine afforded less effective protection against a heterologous B.1.1.529 infection than vaccination with the homologous B.1.1.529-repRNA-CoV2S vaccine. Cumulatively, these data confirm the ability of current vaccines based on the ancestral A.1 strain to provide some degree of protection from B.1.1.529 but an updated homologous SARS-CoV-2 vaccine could provide ideal protection from B.1.1.529 infection.

**Figure 3.**
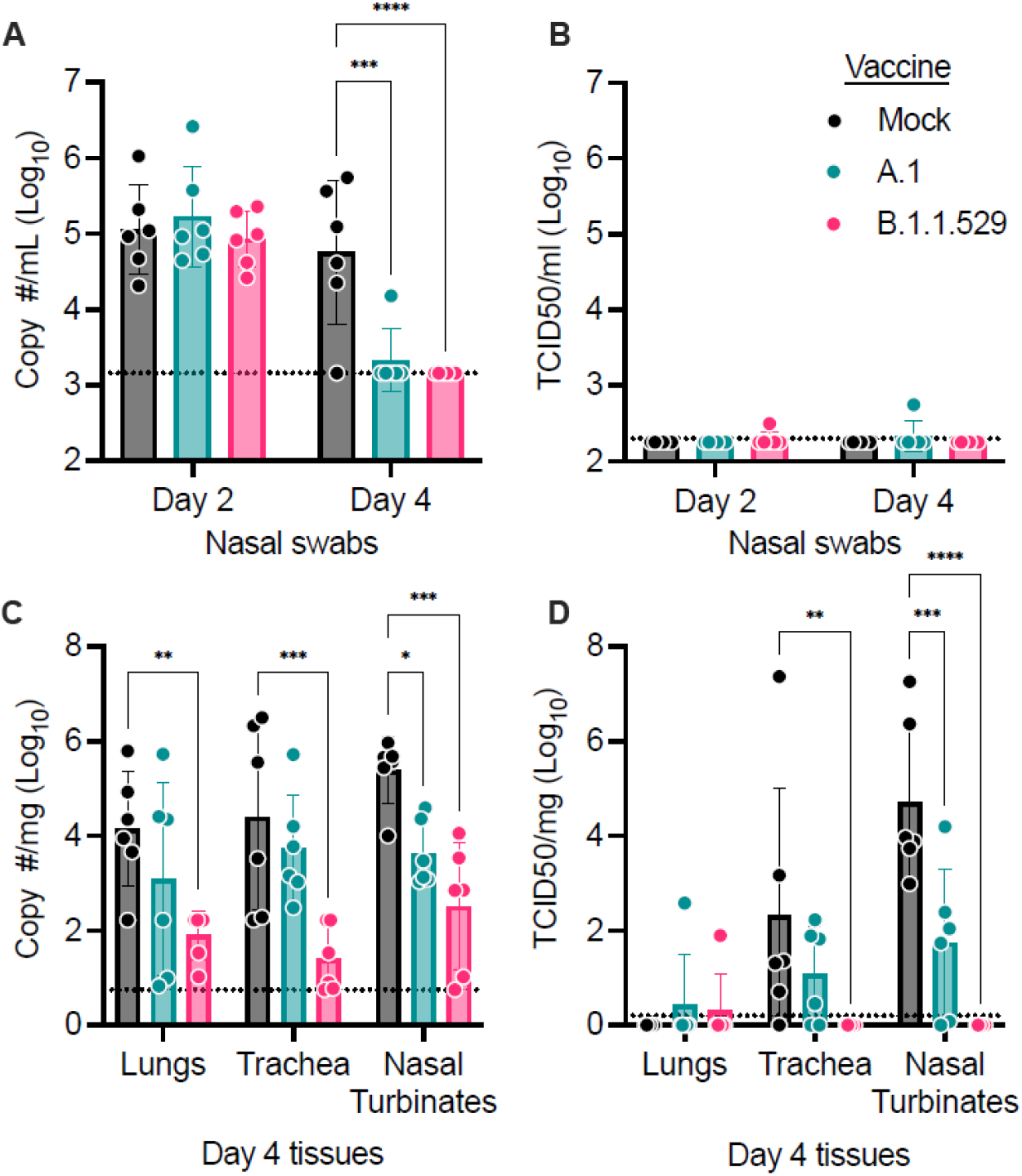
Vaccine protective efficacy against B.1.1.529 infection of Syrian hamsters. Following intranasal infection of vaccinated hamsters with 1000 tissue culture 50% infectious doses (TCID_50_), nasal swabs were collected on days 2 and 4 for evaluation of (**A**) viral RNA load by reverse transcription quantitative polymerase chain reaction (RT-qPCR) or for (**B**) infectious virus by TCID_50_ assay. On day 4, animals were sacrificed and lungs, trachea, as well as nasal turbinates harvested for quantification of (**C**) viral RNA load by RT-qPCR or (**D**) infectious virus by TCID_50_ assay. Indicated statistical comparisons performed using Prism v9. *p<0.05. Comparisons without indicated p-values were non-significant (p>0.05)

Consistent with the low levels of virus in the lungs, histological analysis of formalin-fixed lung sections showed little-to-no lesions, even among mock-vaccinated animals (Fig. 4 and Supplemental Table 1). These data are in contrast to our previous reports on A.1, B.1.1.7 and B.1.351 infection in hamsters that caused significant lung lesions (*17*) and further support the evidence that B.1.1.529 is attenuated in lower respiratory tissue (*20*). Viral antigen was detected in the lungs of 5 of 6 mock-vaccinated animals but in only 1 of 6 and 0 of 6 A.1- or B.1.1.529-repRNA-CoV-2S vaccinated animals, respectively. Hemisections of skulls were evaluated for mucosal inflammation in three anatomically distinct regions: transitional epithelium, respiratory (ciliated) epithelium and olfactory epithelium (Fig. 5 and Supplemental Table 1). Mild to moderate inflammation was observed in the transitional and ciliated epithelium of every mock vaccinated hamster. Similarly, mild to moderate inflammation was observed in transitional and ciliated epithelial regions of all A.1-repRNA-CoV2S vaccinated hamsters. Interestingly, inflammation was also evident in the B.1.1.529-repRNA-CoV2S vaccinated animals, with mild inflammation in both transitional and ciliated epithelium in 5 of 6 vaccinated animals and moderate inflammation in these regions in one hamster. Inflammation was not observed in the olfactory epithelium of any vaccinated or mock-vaccinated hamster (Fig. 5). However, viral antigen was rarely detected in these tissues in the B.1.1.529-repRNA-CoV-2S-vaccinated animals with one hamster exhibiting immunoreactivity in the transitional epithelial compartment and another one exhibiting a focus of immunoreactivity in the ciliated epithelial region (Fig. 5 and Supplemental Table 1). The rare detection of viral antigen in these tissues is consistent with the reduced viral burdens detected in this tissue (Figure 3C & D). In contrast, viral antigen was detected in at least one cellular compartment of all mock-vaccinated animals and 4 of 6 A.1-repRNA-CoV-2S-vaccinated animals had viral antigen in the ciliated epithelial compartment (Fig. 5). The complete histological and IHC findings are provided in supplemental table 1. Cumulatively, these data support the virological data and suggest that the B.1.1.529-repRNA-CoV-2S vaccination provides greater protection against B.1.1.529 challenge than the ancestral A.1 vaccine.

**Figure 4.**
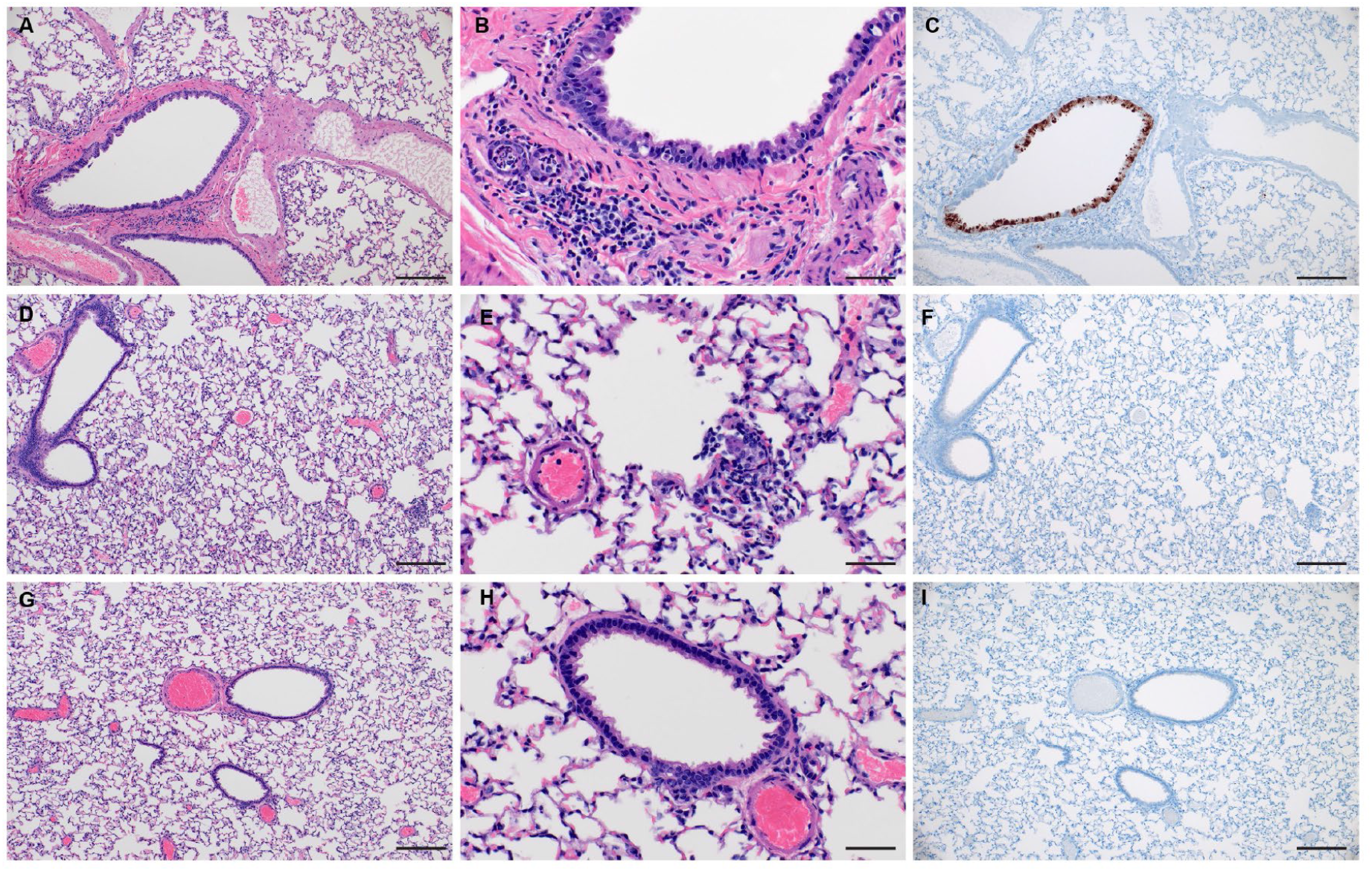
Lung histopathology and immunohistochemistry in vaccinated hamsters. On day 4 PI, lungs were removed and fixed in formalin. Paraffin embedded sections were stained with hematoxylin and eosin (H&E) or with an antibody to detect the SARS-CoV2 N protein (IHC). (A) Section of affected bronchiole with mild bronchiolitis. (B) Higher magnification of bronchiole exhibiting individual epithelial cell necrosis and rare neutrophil infiltration spilling into subjacent submucosal glands. (C) Immunoreactivity is observed primarily in bronhiolar epithelial cells and rarely in alveolar macropaghes (not depicted). (D) Section showing rare, small focus of interstitial pneumonia. (E) Higher magnification of focus of interstitial pneumonia. (F) Immunoreactivity is not observed in association with inflammation. (G) Section of bronchiole and adjacent alveolar spaces lacking any histopathologic lesions. (H) Lack of bronchiolar inflammation at higher magnification. (I) Immunoreactivity is not observed in bronchiolar epithelium or adjacent alveolar spaces. Images shown at 100x (A, D, G, C, F, I, scale bar = 100 μm) or 400x (B, E, H, scale bar = 50 μm).

**Figure 5.**
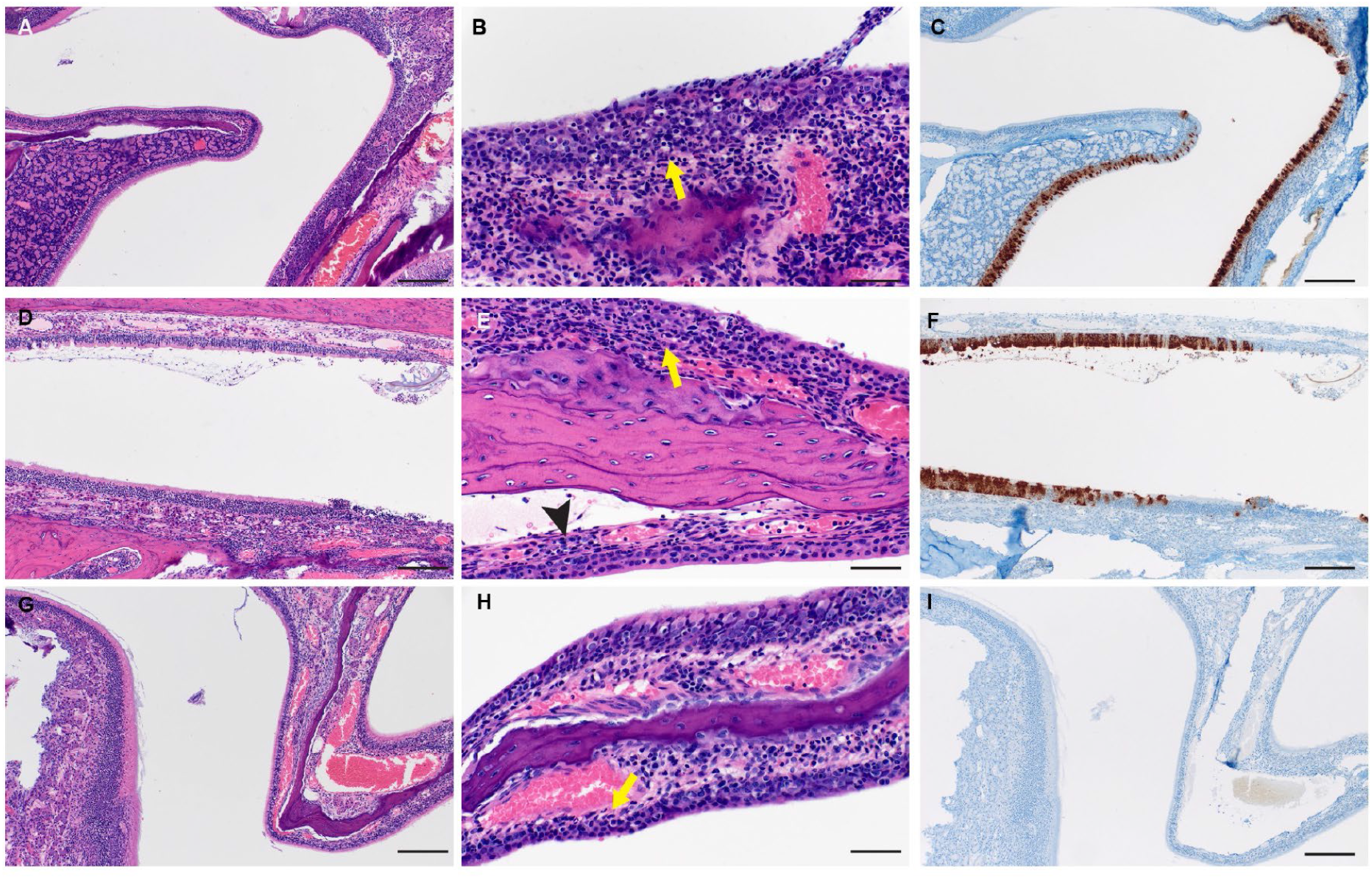
Upper respiratory tract histopathology and IHC in vaccinated hamsters. On day 4 PI, the skull was sectioned and fixed in formalin. Paraffin embedded sections were stained with hematoxylin and eosin (H&E) or with an antibody to detect the SARS-CoV2 N protein (IHC).). (A) Low magnification image showing junction of olfactory epithelium (left) and ciliated epithelium (right) with inflammation within the ciliated epithelium. (B) High magnification of ciliated epithelium with mild neutrophilic influx into the lamina propria (arrow) and individual epithelial cell necrosis in the overlying epithelium. (C) Diffuse immunoreactivity to SARS-CoV-2 is observed in ciliated epithelium with a lack of immunoreactivity in the olfactory epithelium. (D) Low magnification with mild nasal turbinate inflammation and small amounts of luminal exudate. (E) Higher magnification showing mild inflammation in the ciliated epithelium (arrow) and transitional epithelium (arrowhead). (F) Moderate numbers of ciliated epithelial cells exhibiting immunoreactivity to SARS-CoV-2 antigen. (G) Low magnification of olfactory and ciliated epithelial junction with minimal to mild inflammation in the ciliated epithelium. (H) Higher magnification of ciliated epithelum exhibiting rare individual epithelial cell necrosis and influx of low numbers of neutrophils in the lamina propria. (I) Lack of SARS-CoV-2 immunoreactivity in both the ciliated and olfactory epithelium. Images shown at 100x (A, D, G, C, F, I, scale bar = 100 μm) or 400x (B, E, H, scale bar = 50 μm).

## Discussion

The rapid emergence of the B.1.1.529 SARS-CoV-2 VoC has led to a global resurgence of COVID-19 cases, even in heavily vaccinated communities. Multiple reports have shown that immunity conferred by vaccination or recovery from infection with previous SARS-CoV-2 variants provides limited protection against symptomatic infection with B.1.1.529 (*21–23*). Cumulatively, these data suggest that B.1.1.529 can efficiently evade vaccine- or infection-induced immunity driven by previous strains of SARS-CoV-2 (*22, 24*). Our data support this hypothesis as hamsters vaccinated with our vaccine expressing the A.1 spike developed little-to-no neutralizing activity against B.1.1.529 and had inferior protection compared to hamsters vaccinated with the repRNA vaccine expressing the B.1.1.529 spike. Further, in a previous report comparing vaccination with vaccines expressing the spikes of the A.1, B.1.1.7 and B.1.351 SARS-CoV-2 strains and homologous or heterologous SARS-CoV-2 challenge, we found that although neutralizing titers against heterologous strains of SARS-CoV-2 were diminished compared to the homologous strain, protection against viral replication and pathology in the lower respiratory tree remained robust (*17*). These findings suggest sufficient cross-protective immunity to prevent disease, either through non-neutralizing antibody or T-cell activity, is elicited by the A.1, B.1.1.7, and B.1.351 spikes. These data are consistent with reports that A.1 vaccine-induced immunity remained protective against the B.1.1.7, B.1.351 and B.1.617.2 VoCs with diminished protection against B.1.1.529 (*25, 26*).

An important consideration for global public health strategies against current and future emergent VoC is pre-existing immunity from vaccination and/or previous infection. Our data showed that a single booster immunization with a B.1.1.529-targeted vaccine in mice previously vaccinated twice against A.1 failed to develop significant B.1.1.529-specific nAbs whereas mice that received only a single dose of A.1 vaccine were able to mount increasing B.1.1.529-specific bAb antibody responses after two doses of B.1.1.529 vaccine. These results suggest a potential interfering role of the pre-existing immunity and/or antibody levels and are consistent with results observed in human clinical trials testing a heterologous monovalent or bivalent VoC booster vaccines in A.1 pre-immune subjects. In these studies, nAb responses against heterologous virus were not improved following a heterologous booster compared to those who received another A.1 homologous booster (*27, 28*). These data suggest that boosting immune individuals with vaccines specific for an emergent VoC may fail to confer robust immunity against the VoC or could require more than one booster. The mechanisms underlying these findings are not clear. However, like the animals in the study reported here, the human subjects in the cited studies had low to moderate levels of circulating A.1-specific antibody at the time of the boost that may have interfered with interactions between mRNA-produced antigen and B cells or other compartments of the immune system. Alternatively, pre-existing cross-reactive memory B cells could be preferentially recalled, rather than mounting de novo responses via naïve B-cell interactions, as is hypothesized to occur following vaccination in influenza pre-immune individuals (*29*).

In contrast, studies investigating the immunogenicity of a second A.1 booster or 3^rd^ dose in previously vaccinated individuals reported significant increases in serum neutralization activity against not only A.1 but also heterologous B.1.1.529 as well as other VoCs (*21, 22, 30–32*). This effect may be explained by the continued boosting of A.1-specific nAb titers that raises the overall antibody levels and expands rarer cross-neutralizing and/or lower-affinity antibodies, to protective levels. Indeed, a recent study of cases in 10 states as part of the VISION Network, showed that receipt of a third vaccine dose was highly effective at preventing COVID-19-associated hospitalizations during both Delta and Omicron-predominant periods (*33*) However, as A.1-specific immunity wanes, B.1.1.529 immunity is likely to disappear first (*28, 34*). Indeed, breakthrough cases in populations that received a 3^rd^ dose suggest that protection from B.1.1.529 infection after boosting with the current A.1 vaccines is incomplete (*8, 35*), thus prompting the development of updated vaccines that match B.1.1.529. However, our data suggest pre-existing immunity may interfere with protective efficacy of updated vaccines. More data is urgently needed to further characterize and understand the impact of pre-existing immunity on updated vaccines. Lastly, the rapid pace at which B.1.1.529 replaced other circulating SARS-CoV-2 strains (*7*) and evidence that the B.1.1.529 wave has already peaked in countries hit early (*36–38*) suggest that even rapidly produced vaccines based on existing vaccine platforms may be too late to meaningfully impact the course of an emergent VoC (*39*). On the other hand, sub-lineages of B.1.1.529 have already been discovered (*40*) and future VoCs may be descendants of B.1.1.529, making variant-specific vaccines necessary.

Finally, in terms of B.1.1.529 pathogenesis, to our knowledge, our study is the first to report on the histopathology of SARS-CoV-2 in the nasal turbinates of B.1.1.529-infected hamsters. We found mild-to-moderate inflammation in the transitional and ciliated epithelium of either mock- or A.1-repRNA-CoV-2S-vaccinated animals infected with B.1.1.529. The inflammation in these animals was also correlated with the presence of SARS-CoV-2 antigen in the ciliated epithelium. These findings are consistent with previous reports on SARS-CoV-2 infected hamsters that showed inflammation in the turbinates along with the presence of viral antigen (*41*). However, in our B.1.1.529-repRNA-CoV-2S-vaccinated animals, we observed mild-inflammation in the transitional and ciliated epithelium that was not associated with detectable SARS-CoV-2 antigen. This inflammation could be due to host immune responses that are rapidly able to control and clear the inoculated challenge virus. Consistent with other reports on B.1.1.529 infection in hamsters (*14, 18, 19*), we found little to no pathology in the lungs of infected hamsters.

Cumulatively, our data shows that we were able to rapidly synthesize and test a B.1.1.529-targeted vaccine in an *in vivo* hamster model of SARS-CoV-2 infection in response to the discovery of the B.1.1.529 VoC. Our data showed that, compared to an A.1-specific vaccine, a B.1.1.529-targeted vaccine provides superior protection against upper respiratory infection in hamsters and suggest that preexisting immunity may impact the efficacy of variant-specific boosting in immune populations. Importantly, these data indicate that B.1.1.529-targeted immunity will likely be necessary to prevent B.1.1.529 infection and that additional innovations in vaccine technology and design are urgently needed to either overcome or harness pre-existing immunity to drive broadly protective immune responses. Further studies will be needed to develop approaches to drive variant-specific immune responses in pre-immune individuals and to understand how SARS-CoV-2 infection and vaccine boosters impact the breadth of protective immunity to existing and future VoCs.

## Acknowledgements

We thank staff members of the Laboratory of Virology and the Genomics Unit of the Research Technology Branch at Rocky Mountain Laboratories for their efforts in providing a workable SARS-CoV-2 B.1.1.529 stock. Authors also wish to thank the Rocky Mountain Veterinary Branch (NIAID, NIH) and Kevin Draves (UW) for animal care and husbandry.

## Author Contributions

JHE, DWH, HF conceptualized the study. JHE designed the vaccine construct; APK designed the formulation; JHE, APK, KK and JA performed the RNA vaccine production and formulation development and in vitro qualification; JHE, TH, SR performed the mouse experiments; DWH, KMW, CC, SL, DR, ML, RR performed the hamster experiments. SR and TH performed the antibody characterization assays; LH, TYH, ALG, MGJ, ML performed the virus isolation, propagation, and characterization; HF, PB, and DHF provided experimental advice and oversight. DWH and JHE wrote the original draft of the manuscript. DWH, KMW, CC, JA, TH, SL, DR, ML, RR, KK, SR, APK, LH, TYH, ALG, MGJ, PB, DHF, KR, HF, and JHE reviewed and edited the original draft of the manuscript.

## Data and materials availability

All data is available in the manuscript or the supplementary materials.

## Supplementary Materials and Methods

### Biosafety and Ethics

All procedures with infectious SARS-CoV2 were conducted under high biocontainment conditions in accordance with established operating procedures approved by the Rocky Mountain Laboratories (RML) institutional biosafety committee (IBC). Sample inactivation followed IBC approved protocols (Haddock et al., 2021). Animal experiments were approved by the corresponding institutional animal care and use committee and performed by experienced personnel under veterinary oversight. Mice were group-housed, maintained in specific pathogen-free conditions, and entered experiments at 6-8 weeks of age. Hamsters were group-housed in HEPA-filtered cage systems and acclimatized to high containment conditions prior to start of SARS-CoV2 challenge. They were provided with nesting material and food and water ad libitum.

### Viruses and cells

For hamster studies: SARS-CoV-2 variant B.1.1.529 (hCoV-19/USA/GA-EHC-2811C/2021, EPI_ISL_7171744) was obtained from Mehul Suthar, Emory University. Virus stock was sequenced via Illumina-based deep sequencing to confirm identity and exclude contamination. For *in vitro* studies: A.1 lineage SARS-CoV2. used for neutralizing antibody assays, was received from BEI resources. B.1.1.529 lineage SARS-CoV2 was prepared as follows: nasal swab clinical samples recovered from patients undergoing diagnostic testing for the presence of SARS-CoV-2 were collected in viral transport media (VTM) in late December, 2021 by the University of Washington Clinical Virology group and transferred to a biosafety level (BSL)3 laboratory for VTM processing and virus isolation. For virus isolation the VTM was first cleaned by filtering through Corning Costar Spin-X centrifuge tube filter (CLS8160). 0.1 ml of the cleaned VTM was used to infect VeroE6 cells ectopically expressing human Ace2and TMPRSS2 (VeroE6-AT cells; a gift from Dr. Barney Graham, National Institutes of Health, Bethesda MD) in a 48-well plate. Four days later we observed a typical cytopathic effect related to SARS-CoV2 infection. Supernatants were then collected and designated as a passage (P)0 virus stock. The P0 stock was used to produce P1 virus stock, with virus cultures grown in VeroE6/TMPRSS2 cells (JCRB1819). The titer of the P1 stock was measured by standard SARS-CoV-2 plaque assay as described (*16*). An aliquot of the P1 stock was subject to RNA isolation and complete SARS-CoV-2 genome sequencing (RNAseq) using the Swift Biosciences’ SARS-CoV-2 multiplex amplicon sequencing panel (*42*). RNAseq data sets were analyzed by performing phylogenetic comparison across known SARS-CoV-2 sequences present in the Phylogenetic Assignment of Named Global Outbreak Lineages (Pangolin) database (https://github.com/cov-lineages/pangolin) from which we confirmed the isolated virus was the omicron variant. VeroE6-TMPRSS2 (JCRB1819, JCRB Cell Bank, NIBIOHN) cells were cultured at 37C in DMEM supplemented with 10% FBS, 100U/ml of penicillin-streptomycin, and 1mg/ml G418. Baby Hamster Kidney (BHK) cells (ATCC) were cultured at 37C in DMEM supplemented with 10% FBS, and 100U/ml of penicillin-streptomycin.

### Vaccine constructs

A.1-repRNA-CoV2S was previously described (*16,17*). B.1.1.529-repRNA-CoV2 was constructed as follows. GISAID accession EPI_ISL_6699769 was selected for design of 3 overlapping, human codon-optimized, double stranded DNA tiles spanning the entire open reading frame of the spike gene with an additional KV995PP substitution to stabilize the pre-fusion confirmation of spike (also present in the A.1-repRNA-CoV2S construct). Tiles were then synthesized on the BioXP (CodexDNA) and combined with linearized repRNA plasmid backbone in a four-fragment Gibson assembly reaction followed by transformation of e. coli and selection of clones. Sanger sequence-verified plasmid was then scaled prior to linearization by NotI digestion in preparation for transcription and capping as described (*16*). To prepare vaccines for *in vitro* and *in vivo* experiments, RNA was combined with HDT Bio’s stockpiled cationic nanocarrier, Lipid InOrganic Nanoparticle (LION), at a nitrogen-to-phosphate ratio of 15 in a simple 1:1 volume mix and incubated on ice for 30 minutes prior to use.

### Vaccine potency assay

LION/repRNA potency was assayed *in vitro*. Briefly, serial dilutions of LION/repRNA were incubated on a monolayer of BHK cells in a 96-well plate. Twenty-four hours later, cell lysates were added to an ELISA plate coated with anti-SARS-CoV2 Spike (S1 domain) monoclonal antibody. Following a primary incubation and washes, a polyclonal anti-SARS-CoV2 Spike (full-length S) primary antibody was added. Following a secondary incubation and washes, a secondary horse radish peroxidase (HRP)-conjugated antibody was used to detect S-specific binding. Following a final incubation, HRP activity was assayed by TMB/HCL detection and absorbance measured by plate reader (EL_x_808, Bio-Tek Instruments Inc) at 450nm.

### Mouse studies

Six-to-eight-week-old female C57BL/6 mice (Jackson laboratory) received 1μg of each vaccine, as outlined in Figure 2A, via intramuscular injections, in a 50ul volume, on days −52, −24, 0, and 28. Animals were then bled on days 28 and 42, and sera evaluated for neutralizing and binding antibody responses by plaque reduction neutralization test and enzyme linked immunosorbent assay, respectively.

### Plaque reduction neutralization tests (PRNTs)

Two-fold serial dilutions of heat inactivated serum and 600 plaque-forming units (PFU)/ml solution of A.1, or B.1.1.559 viruses were mixed 1:1 in DMEM and incubated for 30 min at 37C. Serum/virus mixtures were added, along with virus only and mock controls, to Vero E6-TMPRSS2 cells (ATCC) in 12-well plates and incubated for 30 min at 37C. Following adsorption, plates were overlayed with a 0.2% agarose DMEM solution supplemented with Penicillin/Streptomycin (Fisher Scientific). Plates were then incubated for 2 (A.1) or 3 (B.1.1.529) days at 37C. Following incubation, 10% formaldehyde (Sigma-Aldrich) in DPBS was added to cells and incubated for 30 minutes at room temperature. Plates were then stained with 1% crystal violet (Sigma-Aldrich) in 20% EtOH (Fisher Scientific). Plaques were enumerated and percent neutralization was calculated relative to the virus-only control.

### Enzyme linked immunosorbent assays (ELISAs)

Antigen-specific IgG responses were detected by ELISA using recombinant A.1 full-length spike, A.1 receptor binding domain (RBD), or B.1.1.529 RBD (Sino Biological). ELISA plates (Corning) were coated with 1 μg/ml antigen or with serial dilutions of purified polyclonal IgG from mice to generate a standard curve in 0.1 M PBS buffer and blocked with 0.2% dry milk-PBS/Tween. Then, in consecutive order, washes in PBS/Tween, serially diluted serum samples, antimouse IgG-HRP (Southern Biotech) and TMB then HCL were added to the plates. Plates were analyzed at 405nm (EL_x_808, Bio-Tek Instruments Inc). Absorbance values from the linear segment of each serum dilution curve was used to interpolate the standard curve and calculate the IgG concentration present in each sample and then fold-change in IgG concentration between days 0 and 28 calculated

### Hamster studies

For hamster studies, Syrian Golden hamsters were purchased from Envigo and were approximately 15-weeks of age at time of vaccination. Hamsters were randomly assigned to study groups and acclimatized for several days prior to vaccination. Hamsters were vaccinated with 20μg of indicated repRNA complexed to LION. RNA was diluted in water and LION diluted in 40% sucrose and 100mM sodium citrate to achieve a theoretical nitrogen:phosphate (N:P) ratio of 15. RNA and LION were allowed to complex for 30 minutes at 4°C. Hamsters were primed with a 50μL intramuscular (IM) injection to each of the hind limbs on day 0. Mock vaccinated hamsters received identical IM immunizations with saline. To monitor antibody responses to vaccination, blood was collected via retroorbital bleeds 27 days after vaccination. Hamsters were monitored daily for appetite, activity and weight loss and no adverse events were observed among the LION/repRNA vaccinated groups. For SARS-CoV2 challenge, hamsters were inoculated with 1000 TCID_50_ indicated SARS-CoV2 variant via 50μL intranasal instillation. Following challenge, hamsters were weighed and monitored daily. Hamsters were orally swabbed on days 2 and 4 post-infection (PI). Swabs were placed in 1mL DMEM without additives. A scheduled necropsy at day 4 PI was performed on all animals to harvest blood and tissues. Studies were performed once.

### Viral RNA quantification

Viral RNA from swabs was isolated using Qiamp RNA mini kit (Qiagen) and viral RNA was isolated from tissues using RNEasy mini kit (Qiagen) according to provided protocols. Viral RNA was quantified by one-step qRT-PCR using QuantiFast Probe PCR reagents (Qiagen) and primers and probes specific for the SARS-CoV2 sub-genomic E RNA as previously described (*43*). For both assays, cycling conditions were as follows: initial hold of 50°C for 10min, initial denaturation of 95°C for 5min, and 40 cycles of 95°C for 15sec followed by 60°C 30sec. SARS-CoV2 RNA standards with known copy number were prepared in house, diluted, and run alongside samples for quantification. The limit of detection was based on the standard curve and defined as the quantity of RNA that would give a Ct value of 40.

### Infectious virus titration

Infectious virus in swabs or tissues was quantified by tissue-culture infectious dose 50 assay (TCID_50_) on Vero cells. Tissues were weighed and homogenized in 1mL DMEM supplemented with 2% FBS and penicillin and streptomycin. Homogenate was clarified of large debris by centrifugation. Samples were then serially 10-fold diluted in DMEM 2% FBS and applied to wells beginning with the 1:10 dilution in triplicate. Cells were incubated for six days before cytopathic effect (CPE) was read. TCID_50_ was determined by the Reed and Muench method (*44*). The limit of detection was defined as at least two wells positive in the 1:10 dilution.

### Histopathology and Immunohistochemistry

At time of necropsy, lungs were dissected and insufflated with 10% neutral buffered formalin. The skull was sectioned and lungs and skull sections submerged in 10% neutral buffered formalin for a minimum of 7 days with 2 changes. Tissues were placed in cassettes and processed with a Sakura VIP-6 Tissue Tek, on a 12-hour automated schedule, using a graded series of ethanol, xylene, and ParaPlast Extra. Prior to staining, embedded tissues were sectioned at 5 μm and dried overnight at 42°C. Specific anti-CoV immunoreactivity was detected using Sino Biological Inc. SARS-CoV/SARS-CoV-2 N antibody (Sino Biological cat#40143-MM05) at a 1:1000 dilution. The secondary antibody was the Vector Laboratories ImPress VR anti-mouse IgG polymer (cat# MP-7422). The tissues were then processed for immunohistochemistry using the Discovery Ultra automated stainer (Ventana Medical Systems) with a ChromoMap DAB kit (Roche Tissue Diagnostics cat#760-159). Sections were scored by a certified pathologist who was blinded to study groups.

### Statistical Analyses

Statistical analyses as described in the figure legends were performed using Prism 8.4.3 (GraphPad).

---

**Supplemental Figure 1:**
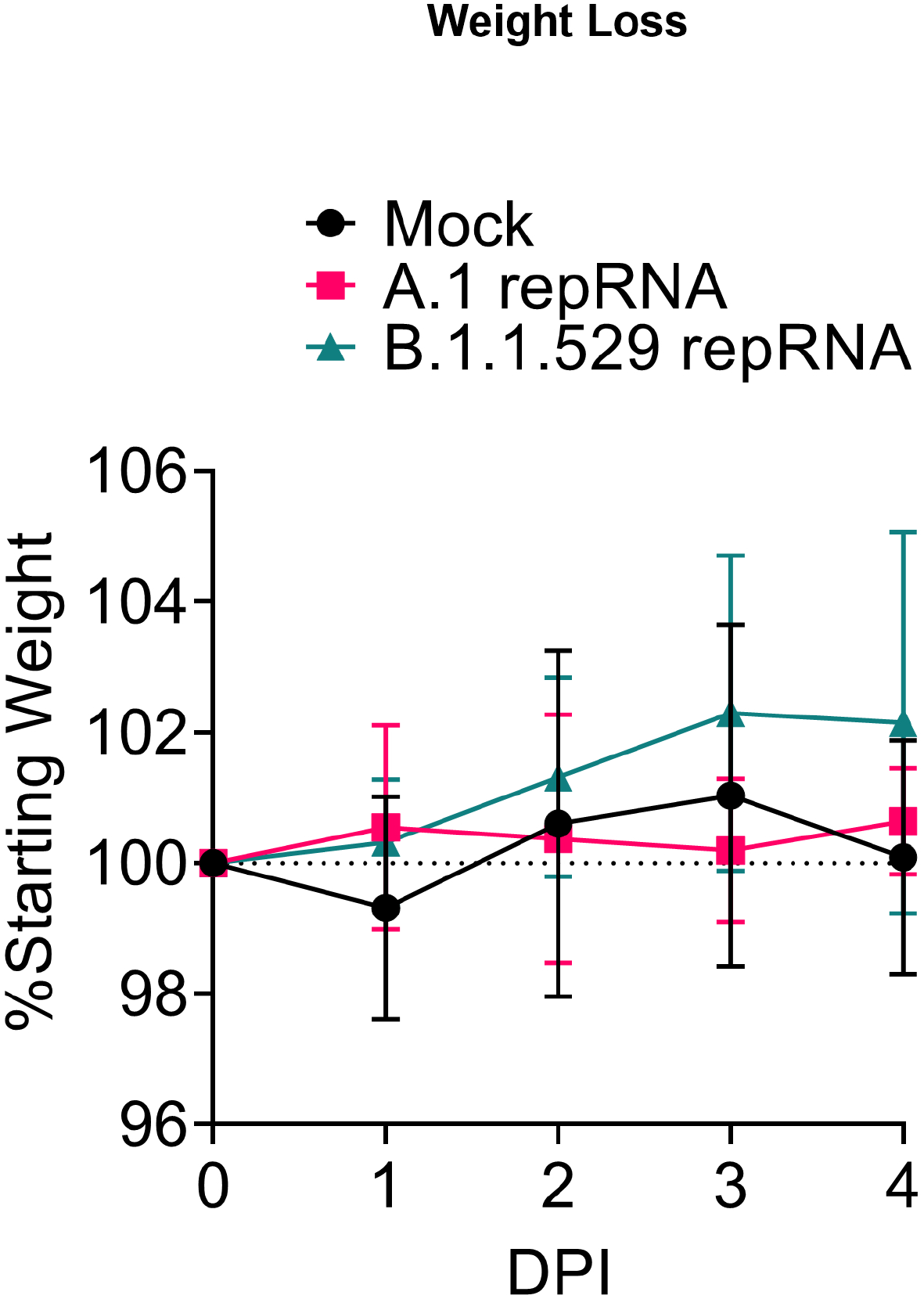
Weight loss in B.1.1,529-infected hamsters. Hamsters vaccinated with indicated repRNAvaccine were challenged with 1000 TCID_50_ of B.1.1.529 via the IN route four weeks after vaccination. Hamsters were weighed daily. Data presented as mean plus standard deviation. N = 6 per group.

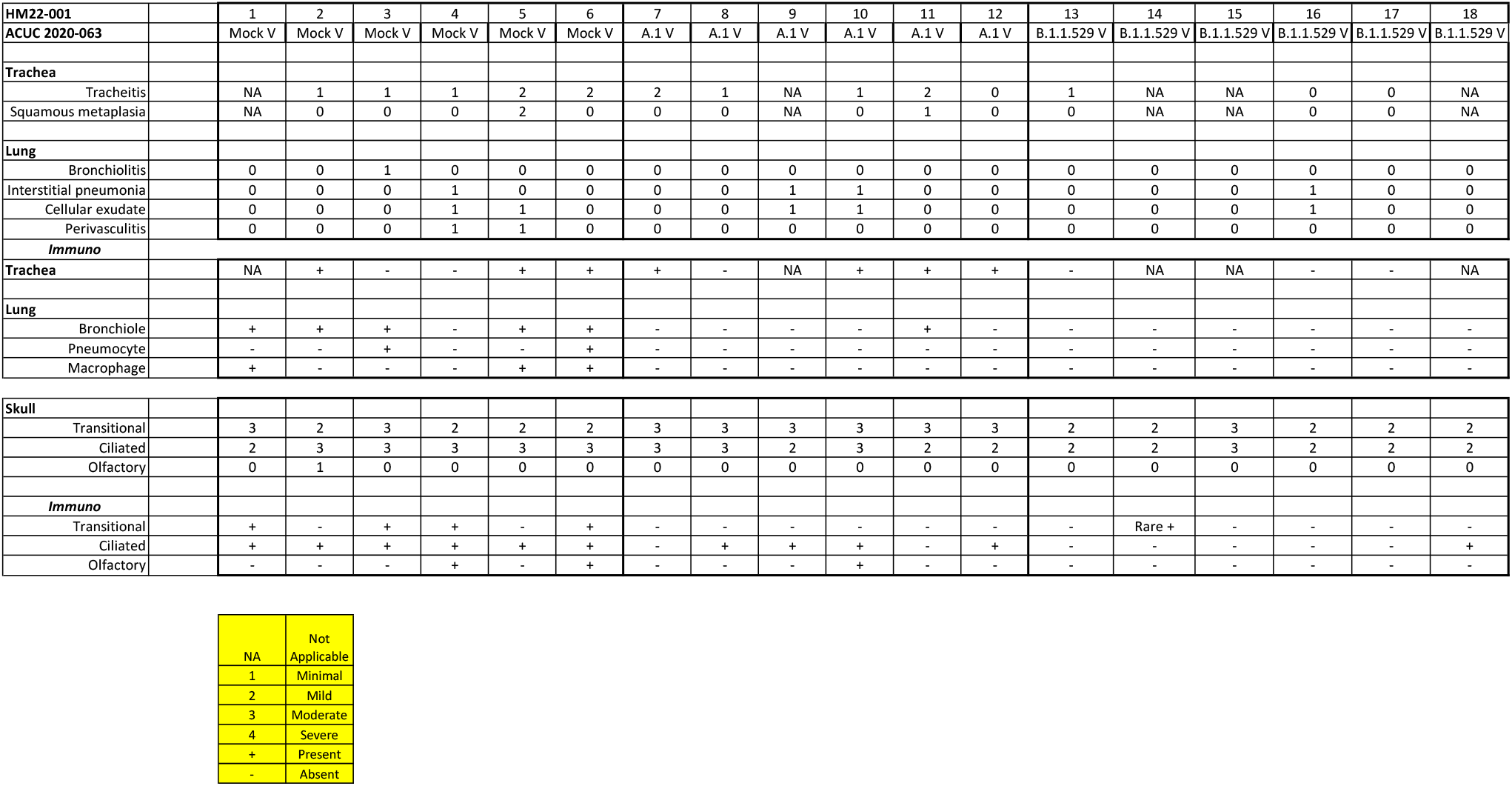

